# QTL mapping of intestinal neutrophil variation and inflammation between threespine stickleback populations reveals links to neurodegenerative disease

**DOI:** 10.1101/757351

**Authors:** Emily A. Beck, Mark C. Currey, Clayton M. Small, William A. Cresko

## Abstract

Host selection is often required to foster beneficial microbial symbionts and suppress deleterious pathogens. In animals, the host immune system is at the center of this relationship. Failed host immune system-microbial interactions can result in a persistent inflammatory response in which the immune system indiscriminately attacks resident microbes, and at times the host cells themselves, leading to diseases such as Ulcerative Colitis, Crohn’s Disease, and Psoriasis. Host genetic variation has been linked to both microbiome diversity and to severity of such inflammatory disease states in humans. However, the microbiome and inflammatory states manifest as quantitative traits, which encompass many genes interacting with one another and the environment. The mechanistic relationships among all of these interacting components are still not clear. Developing natural genetic models of host-microbe interactions is therefore fundamental to understanding the complex genetics of these and other diseases. Threespine stickleback (*Gasterosteus aculeatus*) fish are a tractable model for attacking this problem because of abundant population-level genetic and phenotypic variation in the gut inflammatory response. Previous work in our laboratory identified genetically divergent stickleback populations exhibiting differences in intestinal neutrophil activity. We took advantage of this diversity to genetically map variation in an emblematic element of gut inflammation – intestinal neutrophil recruitment – using an F2-intercross mapping framework. We identified three regions of the genome associated with increased intestinal inflammation containing several promising candidate genes. Within these regions we found candidates in the Coagulation/Complement System, NFkB and MAPK pathways along with several genes associated with neurodegenerative diseases commonly accompanying intestinal inflammation as a secondary symptom. These findings highlight the utility of using naturally genetically diverse ‘evolutionary mutant models’ such as threespine stickleback to better understand interactions among host genetic diversity and microbiome variation in health and disease states.

## Introduction

Animals harbor an array of microbes on and in their bodies which perform essential functions that are fundamental to host health (Fraune and Bosch 2010; Relman 2012; McFall-Ngai et al. 2013). Maintaining appropriate host-microbe interactions by facilitating the presence of symbionts and removing pathogens is therefore vital to sustaining health (Bates et al 2006; Blaser and Falkow 2009; Round and Mazmanian 2009; Chung et al. 2012; Jostins et al. 2012). Interactions between the host immune system and resident microbes are at the center of this relationship (Bates et al 2006; Ley et al. 2008; Blaser and Falkow 2009; Round and Mazmanian 2009; Chung et al. 2012; Jostins et al. 2012; Relman 2012; McFall-Ngai et al. 2013). The immune system can promote beneficial microbes that increase host fitness, and failed interactions can result in a persistent inflammatory response, with the immune system chronically responding negatively to resident microbes. This in turn results in diseases such as Ulcerative Colitis and Crohn’s Disease (Eckburg and Relman 2007; Emilsson et al. 2008; Graham and Xavier 2013).

The relationship between host immune system and resident microbes is complex. Some microbes cause disease states only in specific host genetic backgrounds or in the presence of other microbes (Casadevall and Pirofski 2000). For example, important work in humans has revealed a strong influence of genetic variation on health outcomes particularly in the context of additional microbiome variation (Dethlefsen et al. 2007; Manolio et al. 2009; Ko et al. 2009; Torkamani et al. 2012; Goodrich et al. 2014). In addition, these host-microbe interactions can be mediated by internal environmental conditions such as stress physiology (Lupp et al. 2007; Alverdy and Luo 2017; Mackenzie et al. 2017) and external conditions such as diet (Hildebrandt et al 2009; Albenberg and Wu 2014; Voreades et al. 2014; Singh et al. 2017). As such, host genetic variation and associated microbiomes can productively be considered quantitative traits.

What is needed are studies that can link quantifiable microbe-induced differences in immune response to host genomic loci and genetic variants. One way to quantify the inflammatory response is through assessment of neutrophils, specialized white blood cells that are recruited during an inflammatory response (Bradley et al. 1982; Renshaw et al. 2006; Kumar and Sharma 2010; Mantovini et al. 2011; Kolaczkowska and Kubes 2013). These cells exist throughout the body and are recruited from the blood stream to sites of inflammation, including the gut (Borregard 2010; Fournier and Parkos 2012; Wera et al. 2016). While intestinal neutrophil recruitment often occurs due to the presence of pathogens, it can also occur chronically due to aberrant interactions between the immune system and the gut microbiota (Foell et al. 2002; Wera et al 2016).

Genomic regions that underlie these complex inflammatory phenotypes associated with neutrophil variation can be identified using genetic mapping in model organisms through the use of mutational screens (Musani et al. 2006; Hillhouse et al 2011; Leach et al. 2012; Uddin et al. 2011; Chen et al. 2016; Barry et al. 2018). Because of the complex interplay of genetics, microbes and environment, it is also essential to develop outbred mutant models tractable for genetic mapping of *natural* genetic variants influencing complex phenotypes such as inflammation (Albertson et al. 2009). Here, we use the threespine stickleback fish (*Gasterosteus aculeatus*) as such an outbred ‘evolutionary mutant model’ (Albertson et al. 2009) to study just such complex disease traits.

This small teleost fish is found throughout the arctic in a wide range of environments including freshwater and oceanic habitats, resulting in exceptional degrees of within -and among- population genetic and phenotypic variation for countless traits (Bell and Foster 1994; Colosimo et al. 2004; Cresko et al. 2004, 2007; Hohenlohe et al. 2010; Glazer et al. 2015). Notably, there are multiple high quality genome assemblies from disparate populations (Jones et al. 2012; Peichel et al. 2017) and the large clutch sizes of stickleback provide ample family sizes for QTL mapping (Colosimo et al. 2004; Cresko et al. 2004; Kimmel et al. 2012; Miller et al. 2014; Glazer et al. 2015; Greenwood et al. 2015; Peichel and Marques 2017). By using threespine stickleback lines originating from genetically diverse populations with distinct ecological and evolutionary histories we are able to map natural genetic variants thus allowing us to identify the types of variants likely underlying this complex phenotype in the human population (Albertson et al. 2009).

Previous work in our laboratory described phenotypic variation between freshwater and oceanic ecotype inflammatory responses, with oceanic individuals responding more robustly to the presence of microbes measured by an increase in intestinal neutrophil accumulation and changes in gene expression (Milligan-Myhre et al. 2016; Small et al. 2017). These findings identified a potential role of host genetic variation on differences in intestinal inflammation and the response to the presence of microbes across populations. We set out to map natural genetic variants associated with differences in intestinal neutrophil density using an F2-intercross genetic mapping study in threespine stickleback. We used these data to identify genomic regions that, when combined with previously published gene expression data from juvenile guts in the parental populations (Small et al. 2017), identified a concordant list of candidate genes involved in host immunity. Surprisingly, we also found several genes with characterized functions in neurodegenerative diseases known to include intestinal inflammation as a secondary symptom. These findings have broader impacts in elucidating roles of natural genetic variation in chronic intestinal inflammation and provide further evidence of a strong link between intestinal health and systemic inflammation.

## Methods

### Husbandry and Experimental Design

We generated an F2 mapping cross of threespine stickleback derived from wild caught Alaskan populations previously maintained in the laboratory for at least ten generations. An F1 line was generated by *in vitro* fertilization of parents derived from two distinct Alaskan populations including a male from the freshwater population Boot Lake (N 61.7167, W 149.1167) and a female from the anadromous population Rabbit Slough (N 61.5595, W149.2583). An F2 family (n = 64) was produced intercrossing F1 siblings. Fertilized eggs were incubated overnight in one-micron filter sterilized antibiotic embryo media containing 100 mg/mL Ampicillin, 50 mg/mL Kanamycin, and 8 mg/mL Amphotericin (Milligan-Myhre et al. 2016; Small et al. 2017). The next day embryos were surface sterilized using 0.003% bleach solution and 0.2% Polyvinylpyrolidone-iodine (PVP-I) solution (Western Chemical Inc.) following protocols described by Small et al. 2017. Fish were raised in sterile stickleback embryo media until 9 days post fertilization (dpf) when they were moved to a 9.5 liter tank exposing them to the microbes present in the Cresko Lab fish facility. At this time, fish were housed under “summer” conditions of 16 hours of daylight and 8 hours of night where they were fed 2 mL of hatched brine shrimp naupli and fry food (Ziegler AP100 larval food) designed to mimic the diet of wild stickleback. Water temperature was maintained at 20° C with salinity at 4 parts per thousand (PPT) on a recirculating system. At 14 dpf, fish were euthanized with MS222, following IACUC approved methods described by Cresko et al. 2004. Fish were imaged for standard length measurements, tail-clipped, and fixed overnight in 4% paraformaldehyde (PFA) at room temperature then moved to 4°C for long term storage.

### MPO Staining and Phenotyping

Whole fish were stained using the Sigma Myeloperoxidase kit (Sigma, 390A-1KT, St Louis, MO, USA), which preferentially stains neutrophils. Fish were then embedded in paraffin, and their bodies were cross-sectioned in 7-micron sections from just posterior to the gill rakers to the urogenital opening. Every 10^th^ section beginning posterior to the gill rakers was imaged. Neutrophils were counted twice in each imaged section. If the counts did not agree, the section was counted a third time. Average neutrophils per section were then calculated and tested for association with standard length and sex using R v3.4.1 (R Core Team 2017).

### Statistical Analysis of Phenotypic Variation

Intestinal neutrophil abundance in 14 days post fertilization (dpf) F2s was distributed roughly normally, with full siblings ranging from less than one neutrophil per section on average to over seven neutrophils per section (Figure 1). This variation was shown to be independent of sex (t = 1.35; *P* = 0.18) (Figure S1), as males and females exhibited similar distributions of average intestinal neutrophils. However, neutrophil abundance was shown to be correlated with standard length, with larger fish exhibiting higher average neutrophil density (R^2^= 0.14; *P* = 0.0013) (Figure S2). To account for size as a covariate, we calculated residuals from the regression of neutrophil density and standard length, to be included as trait values in our QTL mapping.

**Figure 1.**
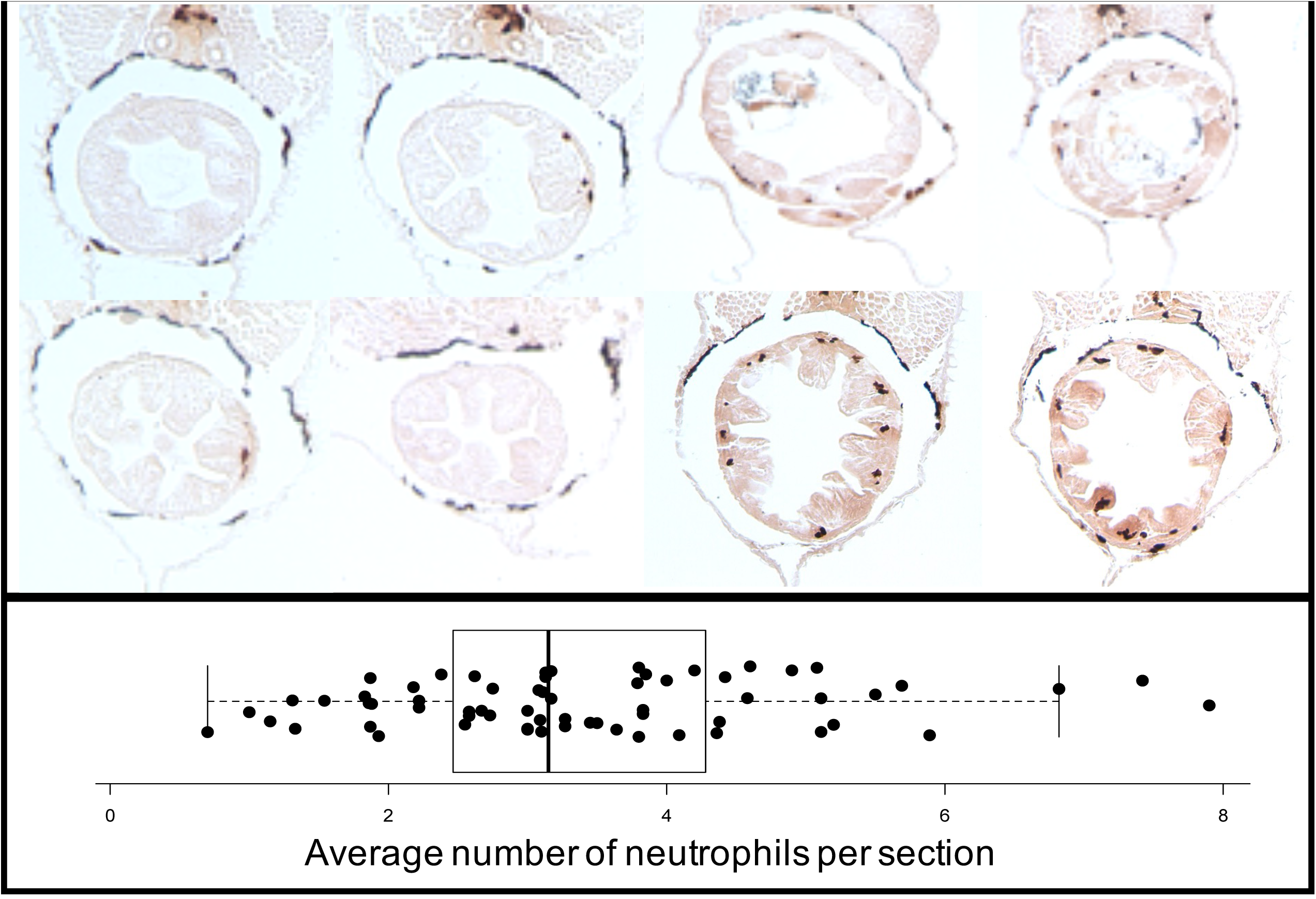
Phenotypic Variation of Intestinal Neutrophil Recruitment. (**A**) Gut sections of a 14 dpf fish with a low average neutrophil count per section. (**B**) Gut sections of a 14 dpf fish with a high average neutrophil count per section. (**C**) F2 Phenotypic distribution of variation of intestinal neutrophil recruitment.

### DNA Isolation and Sex Determination

Tail samples were flash frozen in liquid nitrogen and stored at −80°C. DNA was extracted using the Qiagen DNeasy Blood and Tissue Kit (Qiagen, Valencia, CA, USA). DNA was then quantified using the Qubit fluorometer broad range kit. Individual sex phenotypes were determined using PCR amplification of a sex specific region of the genome using the GA1 primer pair (Griffiths 2000), and males were identified by the presence of the male-specific amplicon.

### Genotyping of Parents and Progeny

Genomic DNA from each F1 parent and F2 offspring was standardized to 10 ng/uL and digested with the endonuclease SbfI-HF (NEB), and RAD-seq libraries were generated using protocols established by the Cresko Laboratory (Hohenlohe et al. 2010; Baird et al 2008; Etter et al 2011). In some progeny samples, DNA concentrations fell below the 10 ng/uL threshold, but all samples with at least 100 ng of DNA were used. Uniquely barcoded samples were then sequenced in one lane on the Illumina HiSeq 4000 to obtain single end 150 bp reads. To improve coverage, the lane was re-run through a second round of sequencing on the HiSeq 4000. Raw sequence data was demulitplexed by barcode and filtered using the process_radtags program in the Stacks suite v1.48 (Catchen et al. 2011; Catchen et al. 2013). Together these sequencing lanes yielded 799,824,397 reads with 708,390,956 reads retained, averaging 3,873,856 reads retained per individual. Reads were then aligned using GSNAP (Wu and Watanabe 2005) to the stickleback reference genome from Ensembl (version 80), allowing for seven maximum mismatches and a terminal threshold of ten. Genotypes were called using the ref_map pipeline of the Stacks suite.

### QTL Mapping

Genotype calls were concatenated in a VCF file generated using the populations package in Stacks (Catchen et al. 2011; Catchen et al. 2013). Filtering was then performed using VCFtools (Danecek et al. 2011). For a marker to be included in the analysis, it was required to have a genotype call in at least 50% of the progeny. MapMaker files were then generated using custom scripts including 18,394 informative markers. QTL mapping was then performed using the r/QTL *scanone* function with Haley-Knott regression (Broman and Sen 2009; Broman et al. 2003). To include growth rate as a covariate, QTL mapping was also performed on residuals calculated from a regression of neutrophils per section on standard length. Genomic regions were further analyzed using the raw neutrophil count data if QTLs were preserved in both the raw and residual data analyses. To account for potential false positives, r/QTL was re-run with phenotypes randomly assigned to genotypes.

### Functional Assignments of Associated SNPs

To identify differentially expressed genes from the parental populations we used RNA-seq differentially expression analysis data from 14 dpf threespine stickleback guts obtained from Small et al. 2017. To assign function to genes within each of our associated SNP boundaries, we used GeneCard v4.8.1 Build8.

### Data Availability

All supplemental files are available on the GSA FigShare portal. Sequencing data are publicly available in the Sequencing Read Archive (SRA) under project ID: SUB879235.

## Results

### Several genomic regions associated with increased intestinal neutrophil density independent of stickleback size

We identified 18,394 informative SNPs to be used for QTL mapping. Using raw intestinal neutrophil densities (number of neutrophils per section), we detected 13 linkage groups (LG) with QTLs associated with variation in neutrophil density (LOD > 3) (Figure 2; Table 1). These included LG1, LG2, LG3, LG4, LG7, LG8, LG9, LG11, LG12, LG13, LG14, LG16, and LG19. To disentangle these findings from standard length and subsequently avoid mapping genes associated with developmental rate or growth, we mapped the residuals of neutrophil number included in a linear model with standard length as a covariate (Figure 2; Table 1). The number of regions of the genome likely associated with intestinal neutrophil recruitment was greatly reduced when mapping the residuals. QTLs retained included pile-ups on LG1, a single SNP on LG4, narrowed regions on LG3 and LG8, and a single pileup on LG12.

**Table 1.**
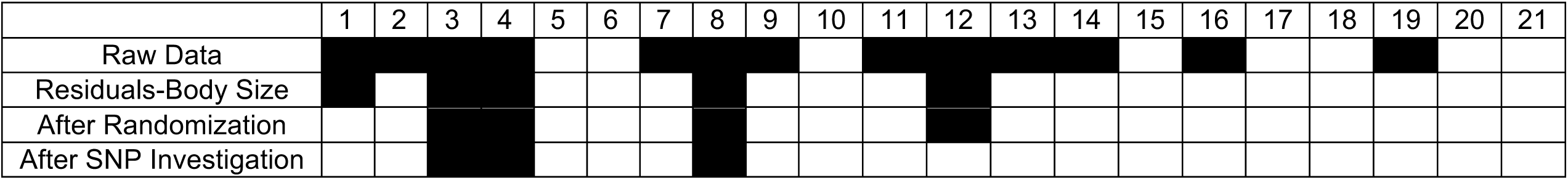
QTLs identified in each Linkage Group (1-21) at each filter step.

**Figure 2.**
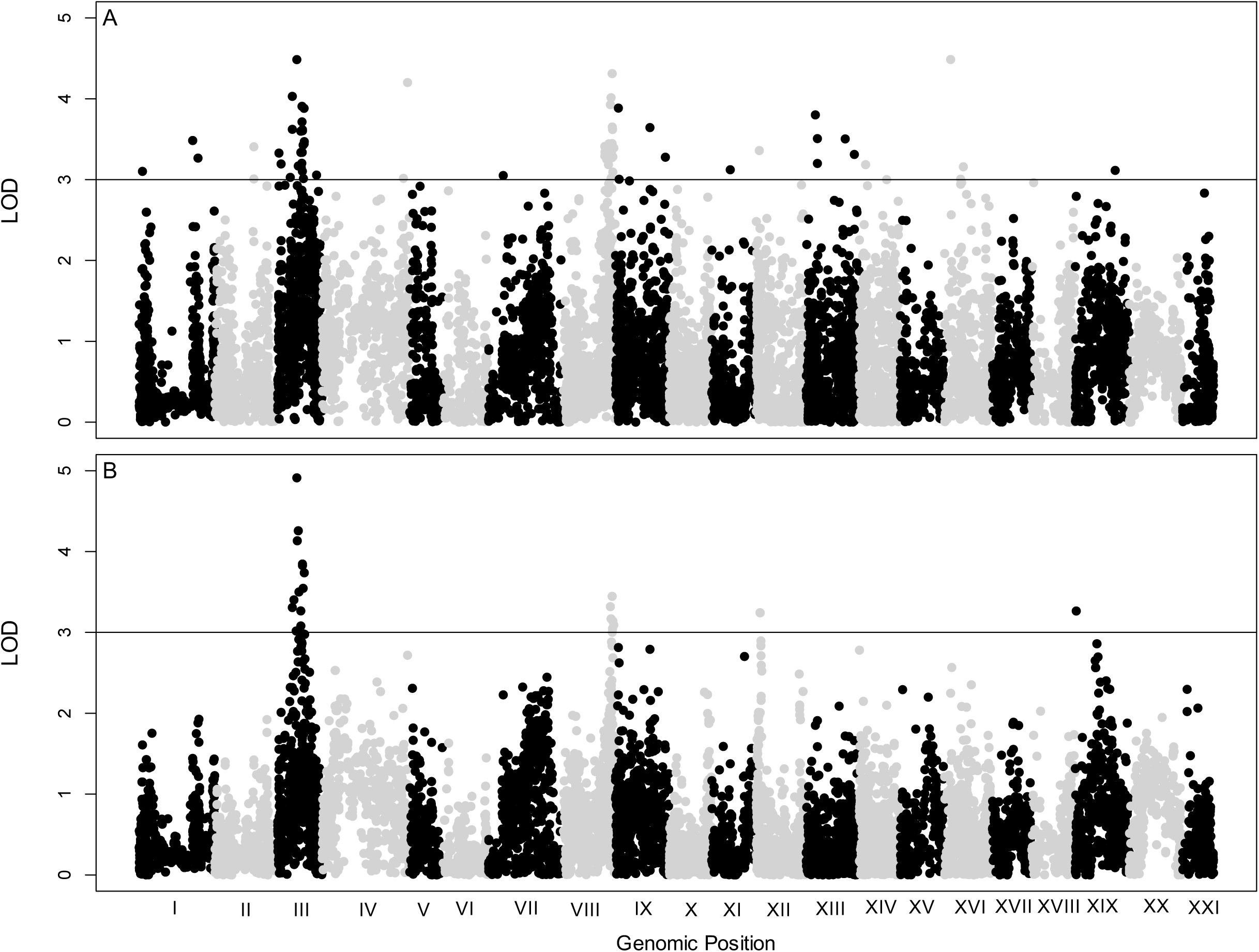
QTL Maps of Average Neutrophil Counts. (**A**) Raw neutrophil count data (**B**) Residual data including growth rate as a covariate. Linkage Groups alternate in color Black/Grey. Horizontal line indicates a LOD cutoff of 3.

In some instances, low coverage or missing genotypes can falsely inflate LOD scores of individual SNPS generating false positives. To evaluate our data for false positives, we randomized the raw phenotype data and the residuals with respect to the F2 genotypes and reran r/QTL. This method is similar to large scale permutation tests used to designate a significant alpha value, but can also indicate regions where low coverage in specific SNPs are generating false positives. In our original runs, high-LOD regions were concentrated in pileups (Figure 2), indicating the type of associative signal expected from linkage mapping. This is in stark contrast to our randomized dataset, with high-LOD SNPs scattered sparsely across the genome (Figure S3). Most importantly, analysis of the randomized datasets revealed very little overlap with our original runs, providing support that low coverage was not generating false positives (Figure S3). The only overlapping signal between the original and randomized data was the pattern of two pileups on LG1. Requiring further analysis of the SNPs on LG1.

To test if low coverage in a few SNPS could be generating false positives, we assessed phenotype distributions or each genotype in the SNPs with LOD > 3. We found false positives on LG1 as expected and additional false positives on LG12. On LG1, four of the five SNPs with LOD >3, neutrophil densities of homozygous individuals did not significantly differ from heterozygous siblings (t = −0.38; *P* = 0.70; df = 48.4; t = 1.36; *P* = 0.18; df = 49.8; t = 1.50; *P* = 0.14; df = 42; t = −1.74; *P* = 0.09; df = 41.5). In the fifth SNP, however, there initially appeared to be a significant increase in neutrophil density in homozygous siblings (t = 2.57; *P* = 0.015; df = 30.6). Upon further inspection, however, we discovered one of the F2s that exhibited a high neutrophil density did not have sufficient coverage at this locus to reliably call a genotype. We imputed the genotype of this missing individual based on surrounding loci, and determined this SNP was likely heterozygous. Reanalysis using the imputed genotype resulted in a loss of significance, suggesting the signal on LG1 was in fact a false positive (t = 1.81; *P* = 0.08; df = 31.6) (Figure S4). Phenotypic distributions consistent with false positives were also identified on LG12 (Figure S5). These markers showed similar patterns to those on LG1 with heterozygous and homozygous siblings exhibiting similar average neutrophil densities (t = −1.11; *P =* 0.28; df = 34.7, t = −1.11; *P =* 0.28; df = 34.7 and t = −1.30; *P* = 0.20; df = 43.4).

At the remaining loci with LOD > 3 on LG3, LG4, and LG8, all testing suggested true associations of SNPs with differences in neutrophil density. On LG3 we subsampled five SNPs exhibiting the highest LOD scores in the pile-up. Three SNPS exhibited a similar pattern: with homozygotes exhibiting a significant decrease in neutrophil density compared to their heterozygous siblings (Table 2; Figure 3; Figure S6). The fourth SNP exhibited the opposite pattern with homozygous siblings exhibiting a significant increase in neutrophil density compared to heterozygous siblings (Table 2; Figure 3; Figure S6). The fifth SNP was generated from two heterozygous parents and therefore was split into three groups: two homozygouys groups (AA/TT) and a heterozygous group (AT) (Figure S6). At this locus, fish homozygous for the A allele had a significantly higher average neutrophil density than either other genotypic group while individuals with with genotypes AT or TT did not differ significantly (Table 2; Figure 3; Figure S6). On LG4 there were three SNPS encompassing two RAD markers with LOD > 3. At each locus heterozygous siblings averaged a higher neutrophil density than homozygous siblings (Table 2; Figure 3; Figure S6). On LG8 we again subsampled the five SNPS with the highest LOD scores which encompassed four RAD markers. In all cases homozygous siblings had a significantly higher neutrophil density than heterozygous siblings (Table 2; Figure 3; Figure S6).

**Table 2.** Phenotypic Distribution summary by SNP.

| Location | Mean<br>Neutrophil<br>Density<br>Homozygote<br>#1 | Mean<br>Neutrophil<br>Density<br>Heterozygote | Mean<br>Neutrophil<br>Density<br>Homozygote<br>#2 | $t^a$ | $df^b$ | $P$ |
| --- | --- | --- | --- | --- | --- | --- |
| **LG3** |  |  |  |  |  |  |
| SNP1 | 3.04 | 4.52 | NA | -3.15 | 29.01 | 0.004 |
| SNP2 | 3.13 | 4.18 | NA | -2.67 | 49.70 | 0.010 |
| SNP3 | 3.06 | 4.66 | NA | -2.99 | 23.00 | 0.006 |
| SNP4 | 4.18 | 3.13 | NA | 2.72 | 50.06 | 0.009 |
| SNP5 | 3.162723 (TT) | 3.057886 (AT) | 5.147527 (AA) | TT/AT: 0.27049<br>TT/AA: -3.5077<br>AT/AA: -4.2347 | TT/AT: 20.16<br>TT/AA: 20.68<br>AT/AA: 15.08 | TT/AT: 0.79<br>TT/AA: 0.002<br>AT/AA: 0.0007 |
| **LG4** |  |  |  |  |  |  |
| SNP1 | 2.83 | 4.13 | NA | -3.90 | 51.83 | 0.0003 |
| SNP2 | 3.12 | 4.24 | NA | -2.77 | 40.67 | 0.008 |
| SNP3 | 3.12 | 4.24 | NA | -2.77 | 40.67 | 0.008 |
| **LG8** |  |  |  |  |  |  |
| SNP1 | 4.16 | 2.84 | NA | 3.87 | 56.93 | 0.0003 |
| SNP2 | 4.16 | 2.84 | NA | 3.87 | 56.93 | 0.0003 |
| SNP3 | 4.30 | 3.06 | NA | 3.28 | 46.92 | 0.002 |
| SNP4 | 4.20 | 2.78 | NA | 4.02 | 52.98 | 0.0002 |
| SNP5 | 4.07 | 2.96 | NA | 3.09 | 53.90 | 0.003 |

a- Welch Two Sample t-test statistic
b- Degrees of freedom

**Figure 3.**
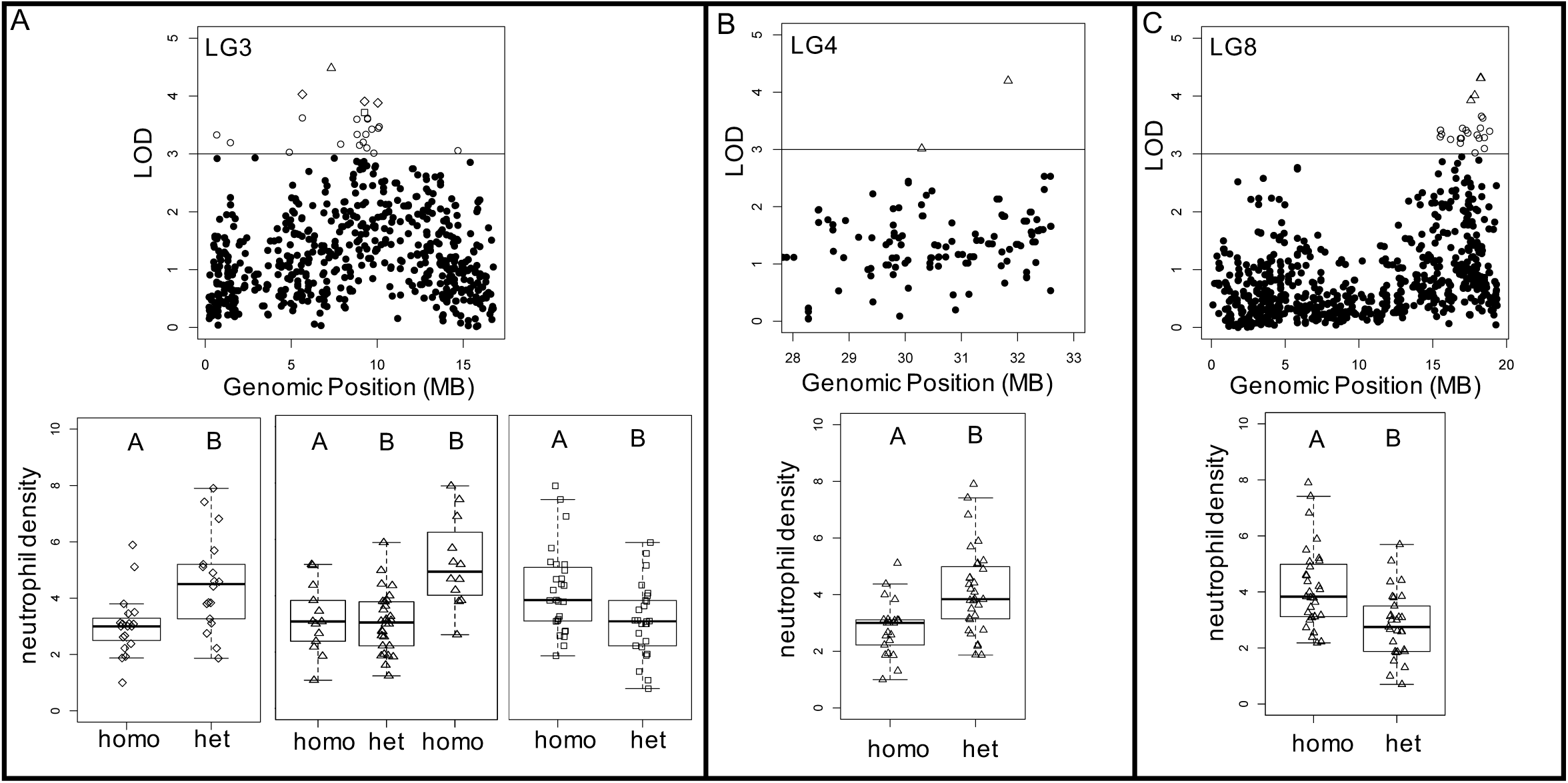
Phenotype Distributions by Genotype. (**A**) Zoomed in view of Manhattan plots of raw QTL data with individual points displaying the highest LOD scores colored based on patterns observed in boxplots. (**B**) Corresponding boxplots to high LOD score SNPs. Relationships between homozygotes and heterozygotes from multiple SNPs are concatenated into single boxplots when they exhibit the same pattern.

To demarcate genomic intervals of interest based on our retained QTLs, we defined boundaries using the outside flanking SNPs LOD < 3 (Table S3). In LG3, we identified 11 SNP clusters LOD > 3, including one marker with a score below the initial threshold of LOD > 3. We chose to include this locus, however, as the LOD score of the residual was one of the highest (LOD = 4.13). In LG4 we identified two genomic regions of interest and in LG8 we identified 17 such regions (Table S3).

### Immune pathways and disease genes associated with increased intestinal neutrophil density

To ascertain biological relevance of each of our QTLs we used the threespine stickleback genome annotation to compile a complete list of candidate genes associated with each genomic interval. Within these boundaries we assembled a complete gene list and identified those with known functions and if available assigned directionality of differential expression from the parental populations (Small et al. 2017). Within these intervals we identified several gene groups on interest including members of the Coagulation/Complement Cascade, Mitogen-Activated-Protein Kinase (MAPK) pathway, Extracellular Signal-Reduced Kinase (ERK), the Nuclear factor kappa-light-chain-enhancer of activated B cells (NfKB) immune pathway, and the maintenance of tight junctions (Table 3).

**Table 3.** Candidate Gene Summary.

| Group | Ensembl ID | Gene | Distance from SNP* (bp) | Immune Pathways | Disease Associations |
| --- | --- | --- | --- | --- | --- |
| LG3 | ENSGACG00000014756 | *f3* | 37,324 | Coagulation Cascade | - |
| LG3 | ENSGACG00000015273 | *map2k2b* | 84,711 | MAPK/ERK | - |
| LG3 | ENSGACG00000015282 | *tcf3a* | 58,000 | MAPK/ERK | - |
| LG3 | ENSGACG00000015301 | *unc13a* | 15,909 | - | ALS |
| LG3 | ENSGACG00000015386 | *epha4b* | 25,437 | MAPK/ERK | - |
| LG3 | ENSGACG00000015719 | *cldn18* | 10,244 | Tight Junctions | - |
| LG3 | ENSGACG00000016028 | *ripk2* | 6,127 | NFkB | - |
| LG3 | ENSGACG00000016066 | *tblxr1b* | 16,774 | - | Autism |
| LG3 | ENSGACG00000016121 | *rgs1* | 1,892 | - | Celiac Disease |
| LG3 | ENSGACG00000016159 | *pik3R3* | 63,634 | MAPK/ERK | - |
| LG3 | ENSGACG00000016189 | *scp2a* | 0 | - | Wheat Allergy |
| LG3 | ENSGACG00000016212 | *angptl3* | 5,349 | MAPK/ERK | - |
| LG3 | ENSGACG00000016323 | *c8b* | 972 | Complement Cascade | - |
| LG3 | ENSGACG00000016338 | *c8a* | 0 | Complement Cascade | - |
| LG4 | ENSGACG00000020015 | *par-4* | 0 | Coagulation Cascade; NFkB | ALS |
| LG8 | ENSGACG00000012681 | *pias4a* | 68,402 | NFkB | - |
| LG8 | ENSGACG00000012686 | *map2k2a* | 58,202 | MAPK/ERK | - |
| LG8 | ENSGACG00000012737 | *tcf3b* | 5,424 | MAPK/ERK | - |
| LG8 | ENSGACG00000013097 | *borcs8* | 19,084 | MAPK/ERK | - |
| LG8 | ENSGACG00000013123 | *ncan* | 0 | MAPK/ERK | - |
| LG8 | ENSGACG00000013599 | *camk4* | 6,656 | MAPK/ERK | - |
| LG8 | ENSGACG00000013618 | *gadd45bb* | 5,900 | MAPK/ERK; NFkB | - |
| LG8 | ENSGACG00000013753 | *gabrb3* | 21,271 | - | Autism |
| LG8 | ENSGACG00000013771 | *gabrg3* | 0 | - | Autism |
| LG8 | ENSGACG00000014026 | *cldn34* | 17,633 | Tight Junctions | - |
| LG8 | ENSGACG00000014029 | *zgc:153311* | 15,831 | Tight Junctions | - |
| LG8 | ENSGACG00000014071 | *nek1* | 80,789 | - | ALS |
| LG8 | ENSGACG00000014199 | *sgcb* | 0 | - | Muscular Dystrophy |

\* indicates closest SNP LOD > 3

The first group of genes were involved in the Coagulation/Complement Cascade, a pathway also enriched for genes differentially expressed between oceanic and freshwater stickleback families (Small et al. 2017). These included *f3, c8a*, and *c8b* on LG3 and *par-4* on LG4 (Table 3). On LG3 one marker was located within the 12^th^ exon of *c8a* while on LG4 a marker was located within intron 1 of *par-4* (Table S4). Interestingly, *par-4* is also differentially expressed in 14 dpf guts of our parental populations, exhibiting significant upregulation in oceanic families compared to freshwater (Small et al. 2017).

The second group included members of the ERK signaling and MAPK pathways. Interestingly, the MAPK Pathway was also enriched for genes sensitive to the presence of microbes in freshwater families (Small et al. 2017). Five of these genes were on LG3: *map2k2b, epha4b, pik3R3, angptl3*, and *tcf3a;* six of these genes were on LG8: *borcs8, ncan, camk4, gadd45bb, map2k2a* and *tcf3b* (Table 3). None of these genes exhibited differential expression patterns (Small et al. 2017) nor were any markers located within the coding region of these genes, but the identification of two co-orthologous pairs (LG3: *map2k2b* and *tcf3a;* LG8: *map2k2a* and *tcf3b*) in addition to the number of genes in this pathway is highly suggestive of a strong association between MAPK/ ERK signaling and intestinal inflammation.

Another pathway associated with several candidate genes was the NfKB immune pathway. This included *ripk2 -* an activator of NFkB – on LG3, *pias4a* and *gadd45bb* on LG8 and *par-4* on LG4 with joint roles in the Coagulation/Complement Cascade and inhibition of the NfKB pathway through direct binding of NFKB1 (Table 3) (Diaz-Meco et al. 1999; Fernandez-Marcos et al 2009; Diaz-Meco and Moscat 2012; Burikhanov et al. 2017). Among these NfKB pathway genes, *par-4* was the only gene differentially expressed in oceanic versus freshwater families, where oceanic families exhibited higher expression (Small et al. 2017).

The last functional group of genes included three genes involved in the formation and maintenance of tight junctions. These included *cldn18* on LG3 and *cldn34* and *zgc:153311* on LG8 (Table 3). Interestingly, *cldn18* and *cldn34* were both upregulated in oceanic compared to freshwater families (Table S4) (Small et al. 2017).

Finally, we identified several gene candidates associated with neurodegenerative diseases that have secondary symptoms relating to intestinal inflammation. This included three genes associated with Amyotrophic Lateral Sclerosis (ALS): *unc13a* on LG3 (Diekstra et al. 2012), *par-4* on LG4 (Xie et al. 2005) and *nek1* on LG8 (Kenna et al. 2016). We also identified three genes associated with Autism Spectrum Disorder (ASD): *tblxr1b* on LG3 and *gabrb3* and *gabrg3* on LG8. Lastly, we identified one disease gene associated with muscular dystrophy, *sgcb*, on LG8.

## Discussion

To our knowledge this is the first study in which an F2 intercross mapping framework has been used to identify genomic loci underlying a complex immune trait. The identification of multiple genomic loci significantly associated with variation in neutrophil density suggests several genes throughout the genome contribute to this complex phenotype. Our results highlight several genomic loci with relatively large effects contributing specifically to intestinal neutrophil recruitment. The persistence of high LOD SNP pile-ups on LG3, LG4, and LG8 after accounting for standard length as a covariate and testing for false positives argue for the biological relevance of the QTLs detected in this analysis. However, the modest size of this single family of F2 progeny (n = 64) means that other genomic regions of small effect would likely have gone undetected. Although other genes of small effect with bearing on the observed variation in intestinal neutrophil activity may have been missed, we can be confident in our identification of several genomic intervals on LG3, LG4, and LG8 with strong associations to intestinal neutrophil activity.

The large number of immune genes identified in the associated genomic intervals is not an unexpected result. The Coagulation/Complement Cascade, MAPK, ERK, and NfkB pathways are all essential immune pathways activate early in development and play roles in the regulation of the inflammatory response (Kurosawa et al. 2000; Markiewski and Lambris 2007; Liu et al. 2007; Dev et al. 2011; Simon et al. 2013; Liu et al. 2017). The identification of a variant within *par-4*, a gene with joint roles in regulation of the Coagulation/Complement Cascade and the NfKB pathway is or particular interest as this gene is known to be upregulated in oceanic populations which exhibit a more robust inflammatory response (Small et al. 2017).

The identification of three co-orthologous pairs was also particularly interesting. In the first pair, *c8a* and *c8b* were both found in the same genomic interval with the associated SNP located within *c8a.* These proteins both function as a part of the Complement Cascade, a pathway enriched for differentially expressed genes between oceanic and freshwater stickleback (Small et al. 2017). Together, these proteins initiate membrane penetration and coordinate the formation and insertion of the membrane attack complex (MAC) into the bilayer to facilitate lysis (Bubeck et al. 2011). The other co-orthologous pairs *map2k2a/map2k2b* and *tcf3a/tcf3b* are also excellent candidates for impacting the inflammatory response as *tcf3* has been shown to play a role in the regulation of B cell maturation and *map2k2* is an activator of MAPK. These pairs are of particular interest, however, as they were found on separate linkage groups with *map2k2b/tcf3a* on LG3 and *map2k2a/tcf3b* on LG8.

The identification of several genes involved in the formation and regulation of tight junctions was also intriguing. Tight junctions are extremely important in regulating intestinal permeability and the intestinal immune response, and have been tied to many intestinal diseases including IBD, Celiac Disease, and Type 1 Diabetes (Visser et al. 2010; Castoldi et al. 2015; Lee et. al 2015; Konig et al 2016). Coinciding with this group were two additional disease genes associated with Celiac Disease and wheat allergies, *rgs1* and *scp2a*, which contained three markers within intronic regions (Table 3; Table S4).

Lastly, the identification of variants within neurodegenerative disease genes provides potential links between intestinal and neurological health. Our findings included variants associated with known disease genes impacting ALS, Autism, and Muscular Dystrophy. All of these diseases are of particular interest to those studying intestinal inflammation, as individuals who exhibit them often report symptoms consistent with colitis and other types on intestinal inflammation at greater rates; and in many cases targeted treatments of the microbiome have been successful in alleviating or slowing progression of symptoms (Nowak et al. 1982; Bellini et al. 2006; Kaneko and Hachiya 2006; Fang 2016; Lo Cascio et al. 2016; Rowin et al. 2017; Hughes et al. 2018; Opazo et al. 2018; Patusco and Ziegler 2018; Spielman et al. 2018; Wright et al. 2018). How the genetics of these complex diseases are tied to intestinal health is still an unresolved problem and requires further mapping of inflammatory phenotypes, but these targets provide a strong starting point to investigate broader implications on intestinal inflammation on systemic health.

## Conclusions

This study provides a strong example of the power of threespine stickleback as a model for mapping natural variants contributing to genetically complex phenotypes relevant to human disease. To expand upon these findings, we can use the stickleback system to map other related immune system phenotypes and expand studies of inflammation to other tissue types. These findings additionally provide potential targets for functional testing using CRISPR-Cas9 genome editing to connect systems-level genetic links between neuro-muscular disorders and intestinal health. Because of the amenability of stickleback for gnotobiotic studies, these genetic approaches will be particularly useful to manipulate both host genes and microbiomes simultaneously to perform functional tests not possible in other organisms. Our findings have broader impacts in elucidating the complex roles of natural genetic variation in chronic intestinal inflammation and provide further evidence of a strong link between intestinal health and systemic health.

## Competing Interests

The authors declare no competing interests.

## Acknowledgements

We would like to thank Susan Bassham and Angel Amores for helpful discussions and comments on the manuscript. We would also like to thank Micaela Burns, Emily Niebergall, Kayla Sharp, and Nathalie Verhoeven for help with imaging and morphometric measurements.

## Author Contributions

The study was designed by E.A.B., C.M.S., and W.A.C. Data Analyses were conducted by E.A.B and M.C.C. The initial manuscript was written by E.A.B with all authors contributing to subsequent versions.

## Funding

This work was supported by the National Institute of General Medical Sciences and the National Center for Research Resources of the National Institutes of Health under award numbers P50GM098911 to (W.A.C, K. Guillemin et al.), R24RR032670 to (W.A.C) and a National Institute of Health NRSA fellowship F32GM122419 to E.A.B.

---

Supplemental Figure 1. Phenotypic Distribution by Sex.

Supplemental Figure 2. Phenotypic Distribution by Standard Length.

Supplemental Figure 3. QTL Maps of Randomized Neutrophil Counts (**A**) Randomized raw neutrophil count data (**B**) Randomized residual data including growth rate as a covariate. Linkage Groups alternate in color Black/Grey. Horizontal line indicates a LOD cutoff of 3.

Supplemental Figure 4. Phenotype Distributions of all analyzed SNPs by Genotype in LG1. Open shapes correspond to boxplots below. Red data points indicate genotype calls missing from the SNP generating a false positive.

Supplemental Figure 5. Phenotype Distributions of all analyzed SNPs by Genotype on LG12.

Supplemental Figure 6. Phenotype Distributions of all analyzed SNPs on (A) LG3 (B) LG4 (C) LG8.

